# A single dose of recombinant VSV-ΔG-spike vaccine provides protection against SARS-CoV-2 challenge

**DOI:** 10.1101/2020.06.18.160655

**Authors:** Yfat Yahalom-Ronen, Hadas Tamir, Sharon Melamed, Boaz Politi, Ohad Shifman, Hagit Achdout, Einat B. Vitner, Ofir Israeli, Elad Milrot, Dana Stein, Inbar Cohen-Gihon, Shlomi Lazar, Hila Gutman, Itai Glinert, Lilach Cherry, Yaron Vagima, Shirley Lazar, Shay Weiss, Amir Ben-Shmuel, Roy Avraham, Reut Puni, Edith Lupu, Elad Bar David, Assa Sittner, Noam Erez, Ran Zichel, Emanuelle Mamroud, Ohad Mazor, Haim Levy, Orly Laskar, Shmuel Yitzhaki, Shmuel C. Shapira, Anat Zvi, Adi Beth-Din, Nir Paran, Tomer Israely

## Abstract

The COVID-19 pandemic caused by SARS-CoV-2 that emerged in December 2019 in China resulted in over 7.8 million infections and over 430,000 deaths worldwide, imposing an urgent need for rapid development of an efficient and cost-effective vaccine, suitable for mass immunization. Here, we generated a replication competent recombinant VSV-ΔG-spike vaccine, in which the glycoprotein of VSV was replaced by the spike protein of the SARS-CoV-2. *In vitro* characterization of the recombinant VSV-ΔG-spike indicated expression and presentation of the spike protein on the viral membrane with antigenic similarity to SARS-CoV-2. A golden Syrian hamster *in vivo* model for COVID-19 was implemented. We show that vaccination of hamsters with recombinant VSV-ΔG-spike results in rapid and potent induction of neutralizing antibodies against SARS-CoV-2. Importantly, single-dose vaccination was able to protect hamsters against SARS-CoV-2 challenge, as demonstrated by the abrogation of body weight loss of the immunized hamsters compared to unvaccinated hamsters. Furthermore, whereas lungs of infected hamsters displayed extensive tissue damage and high viral titers, immunized hamsters’ lungs showed only minor lung pathology, and no viral load. Taken together, we suggest recombinant VSV-ΔG-spike as a safe, efficacious and protective vaccine against SARS-CoV-2 infection.

## Introduction

Severe acute respiratory syndrome coronavirus 2 (SARS-CoV-2) is a member of the *Coronaviridae* family that causes the Coronavirus Disease 2019 (COVID-19) [1–3]. The virus emerged in late 2019 in Wuhan, China and rapidly spread globally. Since then, over 7.8 million cases worldwide were diagnosed, with over 430,000 deaths (as of Jun 10^th^, 2020, covid19.who.int).

SARS-CoV-2 is a single stranded positive sense RNA virus decorated with the spike (S) surface glycoprotein. The S protein is a highly glycosylated type I membrane protein. The homotrimeric organization of the S protein on the viral membrane forms the typical coronaviruses spike structures, [4]. The spike binds with high affinity to the angiotensin-converting enzyme 2 (ACE2) receptor. This binding induces membrane fusion and entry of the SARS-CoV-2 into host cells, thus serving as a target for neutralizing antibodies [5, 6]. The SARS-CoV-2 S protein is composed of two distinct subunits, S1 and S2. The surface unit S1 binds the receptor, whereas the transmembrane unit S2 facilitates viral fusion to cell membranes. The spike is activated by a cleavage at the spike S1/S2 site by host cell proteases [7]. Moreover, it has been recently shown that the SARS-CoV-2 has a newly formed furin cleavage site at the S1/S2 boundary. This novel feature dramatically affects viral entry into Vero E6 and BHK cells [6].

Vesicular Stomatitis Virus (VSV), a member of the *Rhabdoviridae* family, is a non-segmented single-stranded negative sense RNA virus. VSV causes disease in animals, with a broad host range from insects to mammals. However, human VSV infection cases are rare. The VSV genome encodes for five major proteins: matrix protein (M), nucleoprotein (N), large polymerase protein (L), phosphoprotein (P) and glycoprotein (G). The L and P proteins, together with the N, form the transcriptionally active subunit of the virus. The G protein mediates both viral binding and host cell fusion with the endosomal membrane following endocytosis, and cell entry [8].

The recombinant VSV (rVSV) platform was developed by John Rose and Michael Whitt [9, 10]. rVSV was previously used as a vaccine platform for several viral pathogens, including Ebola virus (EBOV), AIDS (HIV) and Crimean-Congo Hemorrhagic Fever (CCHFV) [11, 12]. As a vaccine platform, rVSV harbors several advantages: 1. The virus can be easily propagated and reach high titers. 2. It elicits strong cellular and humoral immunity *in vivo*. 3. Elimination of the VSV-G protein, the major virulence factor of the VSV, attenuates the virus and reduces its reactogenicity. 4. Most of the general population is seronegative for VSV [13].

As the need for a vaccine for SARS-CoV-2 is urgent, more than 90 vaccines are being rapidly developed using a variety of technologies. Among them are RNA and DNA vaccines, viral vectored vaccines, recombinant proteins, live attenuated and inactivated vaccines [14]. Currently, none of these candidates have been approved. Here, we designed an rVSV-based vaccine (rVSV-ΔG-spike), in which the VSV-G protein is replaced with the SARS-CoV-2 S protein, creating a recombinant replicating virus. To this end, we created a cDNA vector encoding the sequence of the N, P, M, and L genes of the VSV genome, and the spike of the SARS-CoV-2, under T7 promoter. We show that the rVSV-ΔG-spike vaccine candidate is expressed by infected cells with the created viral particles robustly expressing the spike protein on the viral membrane. We also demonstrate that rVSV-ΔG-spike is neutralized by SARS-CoV-2 convalescent serum, indicating the antigenic similarity of rVSV-ΔG-spike to the SARS-CoV-2. Moreover, we have implemented a reliable experimental golden Syrian hamster model for COVID-19, enabling rVSV-ΔG-spike vaccine evaluation. A single dose of the rVSV-ΔG-spike vaccine candidate was found to be safe, efficacious, and provides protection against a deleterious SARS-CoV-2 challenge.

## Materials and methods

### Cell lines and viruses

Baby hamster kidney cells (BHK-21, ATCC® CCL-10) and African green monkey kidney clone E6 cells (Vero E6, ATCC® CRL-1586™) were grown in Dulbecco’s modified Eagle’s medium (DMEM) containing 10% Fetal bovine serum (FBS), MEM non-essential amino acids (NEAA), 2mM L-Glutamine, 100Units/ml Penicillin, 0.1mg/ml Streptomycin, 12.5 Units/ml Nystatin (P/S/N) (Biological Industries, Israel). Cells were cultured at 37°C, 5% CO_2_ at 95% air atmosphere.

SARS-CoV-2 (GISAID accession EPI_ISL_406862) was kindly provided by Bundeswehr Institute of Microbiology, Munich, Germany. Virus stocks were propagated (4 passages) and tittered on Vero E6 cells. Handling and working with SARS-CoV-2 virus were conducted in a BSL3 facility in accordance with the biosafety guidelines of the Israel Institute for Biological Research (IIBR). VSV-Indiana (WT-VSV) was kindly provided by Prof. Eran Bacharach (Tel Aviv University).

### Plasmid construction

The pVSV-spike expression plasmid was constructed by PCR amplification of the full length human codon optimized spike gene from pCMV3-SARS-Cov-2 spike expression plasmid (Sino Biological, Cat #VG40588-UT) using the following primers: Forward – atcgatctgtttacgcgtcactATGTTTGTGTTCCTGGTGCTGC; Reverse – atgaagaatctggctagcaggatttgagTCAGGTGTAGTGCAGTTTCACTCC. The amplified PCR product was digested by *MluI* and *NheI* restriction enzymes (NEB), and was ligated into the pVSV-FL+(2) vector (Kerafast), pre-cut by the same enzymes to remove the VSV-G gene. The ligated plasmid was electroporated into DH5α electro-competent cells, and selected by ampicillin resistance.

### Recovery of rVSV-ΔG-spike

BHK-21 cells were infected with Modified Vaccinia Anakara T7 (MVA-T7) virus for 1 hour, followed by co-transfection of five plasmids: the full length rVSV-ΔG-spike (described above), together with the VSV accessory plasmids encoding for VSV-N, P, L and G proteins (Kerafast), all of which under T7 promoter control. The primary transfection was performed using calcium phosphate method. BHK-21 cells were transfected with pCAGGS-VSV-G plasmid to assist in creating passage 1 (P1). 48 hours following primary transfection the supernatant containing the recovered VSV-spike was collected, centrifuged at 1300g X 5min to remove cell debris, and filtered twice using 0.22μM filter to remove residual MVA-T7 virus. pCAGGS-VSV-G transfected BHK-21 cells were then infected with the total amount of the filtered supernatant. 72 hours post-infection the supernatant was collected, centrifuged and used for sequential passaging in Vero E6 cells. rVSV-ΔG-spike was propagated in DMEM containing 5% FBS, MEM NEAA, 2mM L-Glutamine, and P/S/N (Biological Industries, Israel). 15mM D-Trehalose was added to the rVSV-ΔG-spike prior to storage at −80°C.

### Viral titration

Vero E6 cells were seeded in 12-well plates (5×10^5^ cells/well) and grown overnight in DMEM containing 10% FBS, MEM non-essential amino acids, 2nM L-Glutamine, and P/S/N (Biological Industries, Israel). Serial dilutions of rVSV-ΔG-spike were prepared in MEM containing 2% FCS with NEAA, glutamine, and P/S/N, and used to infect Vero E6 monolayers in duplicates or triplicates (200μl/well). Plates were incubated for 1 hour at 37°C to allow viral adsorption. Then, 2ml/well of overlay {MEM containing 2% FBS and 0.4% Tragacanth (Merck, Israel)} was added to each well and plates were incubated at 37°C, 5% CO_2_ for 72 hours. The media was then aspirated and the cells were fixed and stained with 1ml/well of crystal violet solution (Biological Industries, Israel). The number of plaques in each well was determined, and rVSV-ΔG-spike titer was calculated.

### Plaque reduction neutralization test (PRNT)

Vero E6 cells were seeded in 12-well plates as described above. Sera from 12 convalescent COVID-19 patients (approved by the donors for research purposes), rVSV-ΔG-spike vaccinated hamsters, SARS-CoV-2 infected hamsters and mock-infected hamsters were heat-inactivated (at 56°C or 60°C for 30min), then diluted in two-fold serial dilutions (between 1:20 – 1:40,960) in 400μl of MEM containing 2% FCS, NEAA, 2mM L-Glutamine, P/S/N (Biological Industries, Israel), mixed with 400μl of either 300pfu/ml of rVSV-ΔG-spike or SARS-CoV-2, and incubated at 37°C, 5% CO_2_ for 1 hour. Monolayers were washed once with DMEM w/o FBS (for SARS-CoV-2 neutralization only) and 200μl of each serum-virus mixture was added in triplicates to the cells for 1 hour at 37°C. Virus mixture without serum served as control. 2ml/well overlay {MEM containing 2% FBS and 0.4% Tragacanth (sigma, Israel)} were added to each well and plates were incubated at 37°C 5% CO_2_ for 48 (for SARS-CoV-2) or 72 hours (for rVSV-ΔG-spike). Following incubation, overlay was aspirated and the cells were fixed and stained with 1ml of crystal violet solution (Biological industries, Israel). The number of plaques in each well was determined, and the serum dilution that neutralizes 50% of the virions (NT_50_) was calculated using Prism software.

### Immunofluorescence analysis (IFA)

Monolayers of Vero E6 cells were seeded in 8 chamber LabTek (Nunc) (1.5×10^5^ cells/well) and infected with WT-VSV, SARS-CoV-2 or rVSV-ΔG-spike for 24 hours. Cells were then fixed with 3% paraformaldehyde (PFA) in PBS for 20 minutes and permeabilized with 0.5% Triton X-100 for 2 minutes. The fixed cells were blocked with PBS containing 2% FBS and stained with either hyperimmune rabbit serum from intravenous (i.v.) SARS-CoV-2 infected rabbits, hamster sera or COVID-19 convalescent human sera for 1 hour. After washing with PBS, cells were incubated with Alexa Fluor 488 conjugated anti-rabbit, anti-human and anti-hamster antibodies, respectively. Nuclei were stained with DAPI. The cells were visualized by an Axioskop (Zeiss) equipped with a DS-iR1 camera (Nikon) and images were taken using the NIS-elements software (Nikon).

### Ultrastructure analysis

To determine the ultrastructure of the rVSV-ΔG-spike or WT-VSV, transmission electron microscopy (TEM) was performed. Carbon-coated grids were immersed in DDW. Clarified supernatants were absorbed to the grids by a drop-on-grid method (DOG) for 15-20 minutes. Filtered blocking solution of 2% FBS in PBS was added to the grids for 20 minutes. Immuno-gold labeling was performed using polyclonal antibody (pAB RBD SBF40150-T62) directed at the receptor binding domain (RBD) of the spike, followed by washing (x3) with PBS and then labeling with gold-conjugated goat anti-rabbit secondary antibody (Sigma, G3779). The grids were washed with PBS (x3), and DDW (twice). Grids were stained with 1% phosphotungstic acid and examined with a Tecnai T12 TEM {Thermo Fisher Scientific formerly FEI)} operated at 120kV and equipped with a Gatan ES500W Erlangshen camera.

### Real-time RT PCR

RNA was extracted by Viral RNA mini kit (Qiagen, Germany) as per manufacturesr’s instructions. Real-time RT PCR was performed using the SensiFAST™ Probe Lo-ROX one-step kit (Bioline). In each reaction the primers final concentration was 600nM and the probe concentration was 300nM. Primers and probes were designed using the Primer Express Software (Applied Biosystems) and purchased from Integrated DNA Technologies, Inc. Probes were ordered as 6-FAM and ZEN/Iowa Black FQ combination. The primers and probes used:

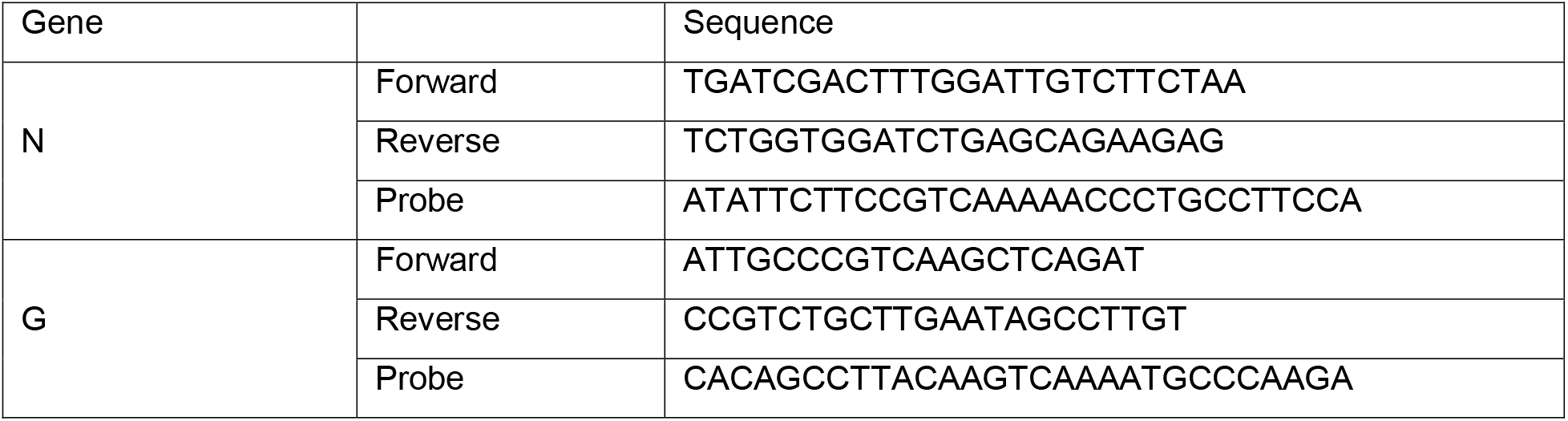
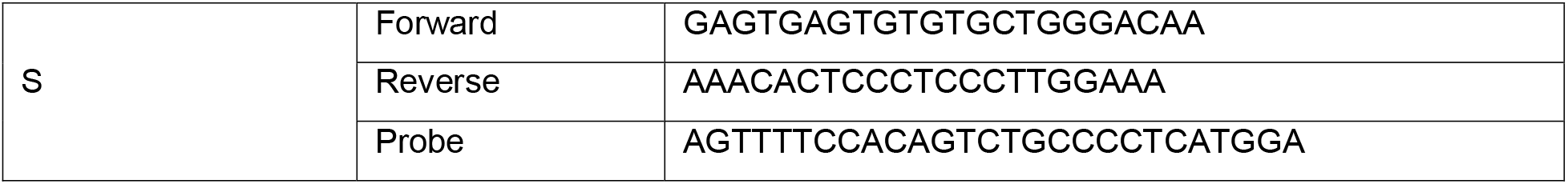

### Whole genome sequencing and data analysis

The SMARTer Pico RNA Kit (Clontech) was used for library preparation. Whole genome sequencing was conducted using the Illumina Miseq platform, with a read length of 60 nucleotides, producing 5,821,469 reads. FastQC [15] was used for quality control of the data. Reads originated from Vero E6 host cells were filtered out using Bowtie 2 [16], resulting in 1,070,483 reads originated from rVSV-ΔG-spike. Mapping of the reads against the rVSV-ΔG-spike was performed using Bowtie 2 followed by variant calling using Samtools [17], both with default parameters, resulting in a 3,178x average coverage and several variants.

### Animal experiments

The animal model for SARS-CoV-2 was established by intranasal instillation (i.n.) of SARS-CoV-2 diluted in PBS supplemented with 2% FBS (PBF) (Biological Industries, Israel) to anesthetized {intraperitoneal (i.p.) Ketamine (160mg/kg) with Xylazine (6mg/kg)} 6-7 weeks old golden Syrian hamsters (60-90 gr., Charles River Laboratories, USA). Animals’ body weight was monitored daily. Five or seven days post infection (dpi), the animals were sacrificed and lungs from infected hamsters were excised and processed for viral load determination, or fixed in 4% PBS-buffered formalin and processed for histopathological analysis as described below.

Vaccination was performed by intramuscular (i.m. 0.05 or 0.1ml/animal) or subcutaneous (s.c. 0.3 ml/animal) injection of rVSV-ΔG-spike to anesthetized golden Syrian hamster (6-7 weeks old, 60-90 gr.). Vaccinated animals’ weight was monitored daily for 11 days post-vaccination. Sera was collected two weeks post-vaccination for titration of SARS-CoV-2 neutralizing antibodies. 20 or 25 days post vaccination, hamsters were anesthetized, challenged i.n. with 5×10^6^ pfu of SARS-CoV-2 and monitored for 11 additional days. All animal experiments involving SARS-CoV-2 were conducted in a BSL3 facility in accordance with the guideline of the Israel Institute for Biological Research (IIBR) animal experiments committee. Protocol numbers: #HM-01-20, HM-02-20, HM-03-20.

### Lung viral load determination

Hamsters’ lungs were harvested at 5 dpi, and stored at −80°C. Lung processing and infectious virus quantitation was performed by plaque assay, as described above.

### Histopathology

For hematoxylin and eosin staining (H&E), the lungs were rapidly dissected and carefully inflated with 0.3ml 4% neutral-buffered paraformaldehyde and further fixed in 4% neutral-buffered paraformaldehyde at room temperature for two weeks, followed by routine processing for paraffin embedding. Coronal, serial sections, 5μm-thick, were performed and selected sections were stained with H&E for light microscopy examination. Images were acquired using Nikon Eclipse 50i Light Microscope (Nikon, Tokyo, Japan).

For Immunolabeling of SARS-CoV-2, sections were deparaffinized and rehydrated through 100% ethanol, 95% ethanol, 70% ethanol and 30% ethanol, washed in distilled water and antigens were retrieved using commercial antigen retrieval solution (Dako, CA, USA). Sections were then permiabilized for 10 minutes (0.2% Triton X-100 in PBS), blocked for 1 hour (10% NGS in PBS containing 0.05% Triton X-100), incubated in 100 μg/ml of rabbit SARS-CoV-2 primary antibody (in-house preparation of rabbit polyclonal anti-RBD) in antibody cocktail solution (50% blocking solution, 0.05% Triton X-100 in PBS) for 24 hours at 4°C.Sections were then washed three times with washing buffer (1% blocking solution in PBS containing 0.05% Triton X-100) and incubated with anti-rabbit Alexa Fluor 488 secondary antibody (Molecular Probes, Burlington, Canada) in antibody cocktail solution for 1 hour at room temperature. Nuclei were stained with DAPI. Following three additional washes, slides were mounted using Fluoromount-G (Southern Biotech, Al, USA) and images were acquired using a Zeiss LSM710 confocal microscope (Zeiss, Oberkochen, Germany).

### Tissue/Air space ratio calculation

Briefly, tissue/air space ratio was determined using Image J free software analysis (particle analysis algorithm). Images of six random regions of interests (ROI) per section were taken at the same magnification (x20). Color threshold parameters were set and remained consistent throughout analysis. Total area values were measured separately for air space and tissue. Ratio of total tissue area to total air space area was calculated for each ROI. Average value of 6 ROIs is presented.

### Statistical analysis

Data was analyzed with GraphPad Prism 5 software (GraphPad software inc.). P<0.05 was considered statistically significant. Morbidity analysis was performed as means of the area under the curve (AUC) in presence of individual weight at the baseline as previously described [18]. Briefly, the AUC was calculated during the entire period, starting from the day 0 (day of challenge), until the end of the experiment. The differences between groups were analyzed using student *t-test*.

## Results

### Generation of rVSV-ΔG-spike

The goal of rapid and efficient mass vaccination calls for the implementation of replicating vaccines, such as VSV-based vaccines [13]. To this purpose, we replaced the open reading frame (ORF) of VSV-G with the full length human codon-optimized spike gene of SARS-CoV-2 (Sino Biological), within the VSV full length expression vector, yielding pVSV-ΔG-spike (Fig. 1A, B). Primary recovery of the VSV-ΔG-spike was performed in BHK-21 cells infected with Modified Vaccinia Anakara T7 (MVA-T7), followed by co-transfection with the rVSV-ΔG-spike, and the VSV accessory plasmids encoding for VSV-N, P, L and G proteins under control of a T7 promoter (Fig. 1C). To further support the entry of the recovered virus during the initial steps, VSV-G was expressed under a strong mammalian promoter. To remove residual MVA-T7 from the recovered virus, two sequential filtration steps were applied followed by infection of Vero E6 cells, as these cells support SARS-CoV-2 spike mediated entry, but not MVA replication. 72 hours post infection, the infected cells underwent syncytia formation, accompanied by significant cytopathic effect (CPE). The supernatant containing the recovered rVSV-ΔG-spike was then collected, centrifuged, and used for further passaging. Thirteen subsequent passages on Vero E6 cells were performed aimed to eliminate the carryover of the VSV-G protein used during the initial recovery steps, and to increase the viral replication efficiency and titer.

**Figure 1.**
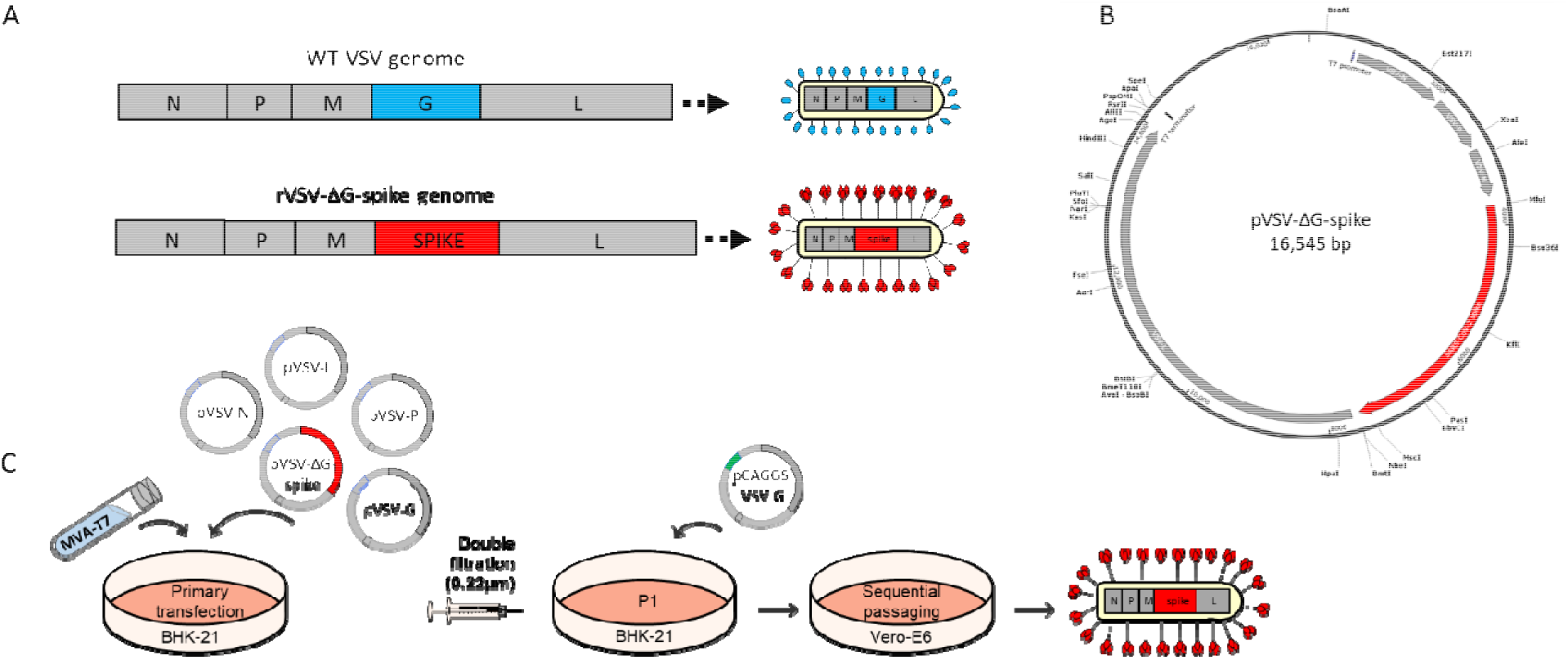
rVSV-ΔG-spike design and generation strategy: (A) A schematic diagram of the genome organization of WT-VSV genome (top diagram) and rVSV-ΔG-spike (bottom diagram). N: Nucleoprotein, P: Phosphoprotein, M: Matrix, L: Large polymerase, G: Glycoprotein, SPIKE: SARS-CoV-2 spike. (B) pVSV-ΔG-spike map; in red, the inserted S gene. (C) Schematic representation of the generation process of creating rVSV-ΔG-spike vaccine. Infection of BHK-21 cells with MVA-T7, followed by co-transfecti with pVSV-ΔG-spike, and VSV-system accessory plasmids; Transfection of BHK-21 cells with pCAGGS-VSV-G, followed by infection with the supernatant of the primary transfection to create P1; serial passaging to create rVSV-ΔG-spike.

### Characterization and analysis of rVSV-ΔG-spike

The above mentioned 13 sequential passages were accompanied by morphological manifestations. The VSV-G gene carryover from the initial transfection was gradually eliminated while maintaining the SARS-CoV-2-S gene, and other VSV genomic RNA-encoded genes (represented by VSV-N gene), as demonstrated by quantitative real time RT-PCR (Fig. 2A).

**Figure 2.**
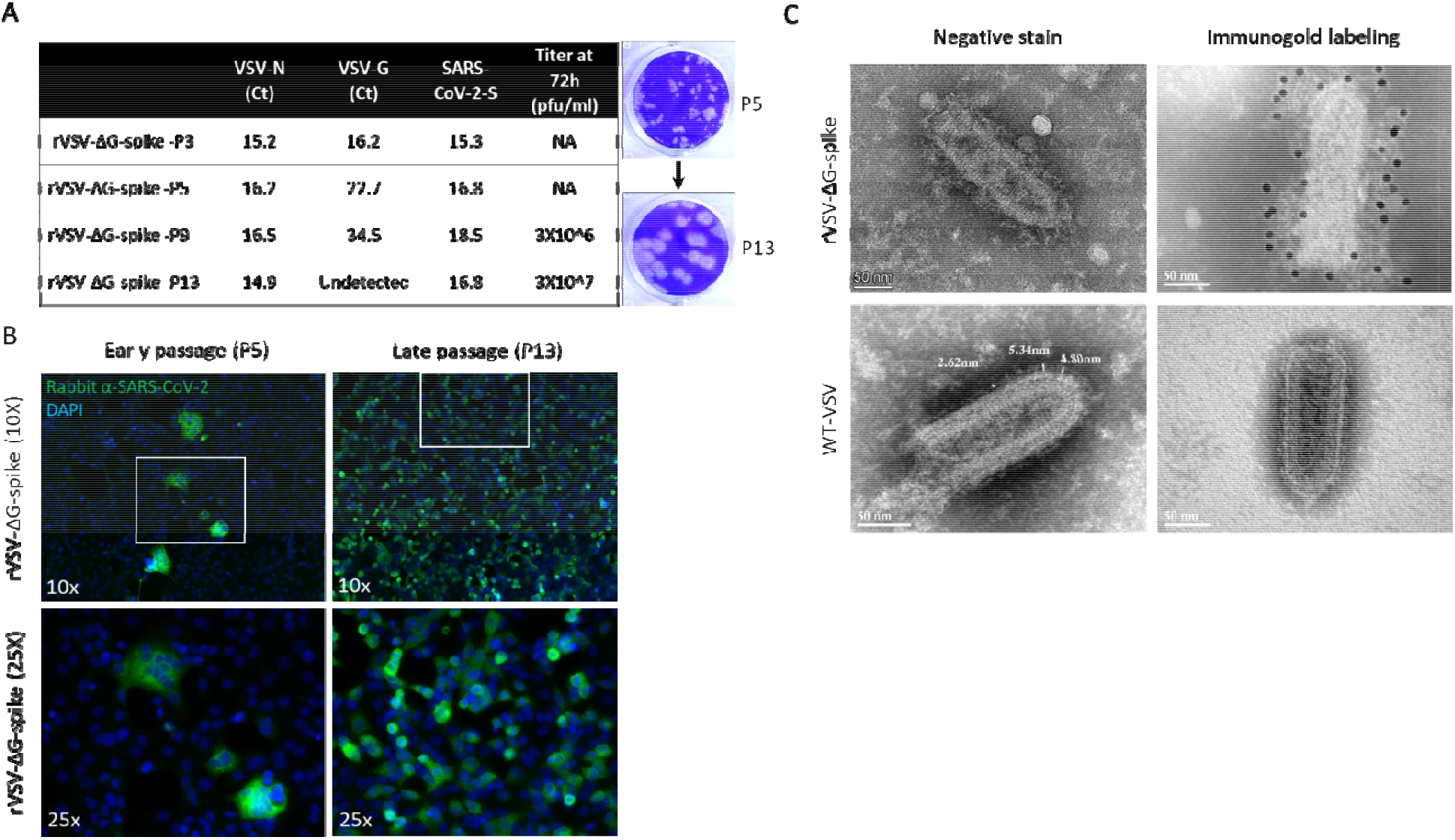
Characterization of rVSV-ΔG-spike: (A) A summary of the genome analysis of several passages of rVSV-ΔG-spike, showing Ct values of VSV-G, VSV-N, and SARS-CoV-2-S indicating the elimination of VSV-G over time, together with increased titer. Also, plaques are formed at late passages. (B) Immunofluorescence images of Vero E6 cells infected with early passage (P5)-rVSV-ΔG-spike, or late passage (P13)-rVSV-ΔG-spike, stained with a SARS-CoV-2 antibody (green) and DAPI for nuclei staining (blue). Top and bottom panels show insets at low (10X) and high magnification (25x), respectively. rVSV-ΔG-spike at P5 forms syncytia, whereas P13 show individual infected cells, with no evidence of syncytia. (C) Transmission electron microscopy of rVSV-ΔG-spike (top panel) compared to WT-VSV. Right panel shows immunogold labeling against RBD.

To assess the growth kinetics of the rVSV-ΔG-spike throughout the sequential passages, rVSV-ΔG-spike titer was determined at several time points by plaque assay. During the initial passages, the effect of rVSV-ΔG-spike was characterized by the formation of large syncytia accompanied by cytopathic effect (CPE), as visualized by cell rounding, detachment, and disruption of the entire monolayer. However, traditional plaques were not observed. Plaque formation was evident only at later stages, correlating with the decrease in syncytia formation, increase in S gene expression and elimination of VSV-G gene. Accordingly, rVSV-ΔG-spike titer was increased during passaging progression, reaching 3×10^7^ pfu/ml, at passage 13 (P13). (Fig. 2A).

To evaluate the expression of rVSV-ΔG-spike, we performed immunofluorescence analysis (IFA) using anti-SARS-CoV-2 antibodies. SARS-CoV-2 spike protein was efficiently expressed in rVSV-ΔG-spike-infected Vero E6 cells (Fig. 2B). Immunostaining of rVSV-ΔG-spike-infected cells displayed syncytia, mostly at the initial passages during the creation of the rVSV-ΔG-spike (P5), whereas during advanced steps (P13) showed a robust expression of the spike protein in individually infected cells.

Transmission electron microscopy (TEM) was performed to analyze the ultrastructure of the rVSV-ΔG-spike. rVSV-ΔG-spike particles maintained the characteristic *Rhabdoviridae* bullet-shape morphology and were decorated with spike protein on the particles’ membrane, already at early stages. Passaging led to increase in the prevalence of the spike structures per single particle (Fig. 2C). Furthermore, the spikes’ lengths were measured, and found to be at the range of 12-20nm, consistent with that of the spike protein on SARS-CoV-2 particles [19]. In contrast, G protein length, ranging from 2-6nm, was observed on WT-VSV particles. Immunolabeling of rVSV-ΔG-spike using polyclonal antibodies directed at the receptor binding domain (RBD) of the spike further validated the presence of spike structures on the rVSV-ΔG-spike viral particle (Fig. 2C).

### Spike mutations acquired during the creation of rVSV-ΔG-spike vaccine candidate

The process of rVSV-ΔG-spike generation by serial passaging was accompanied by sequencing at key passages. While early in the process no significant variations were detected, later passages highlighted the emergence of three mutations (Table 1): (1) A synonymous mutation at position 507 that emerged at later passages, (2) A nonsynonymous mutation R685G - This mutation is located at the multibasic motif of the S1/S2 cleavage site of the spike protein - RRAR, thus creating a RRAG site and (3) A nonsense mutation introducing a stop codon at position 1250 of the spike protein (C1250*), leading to truncation of 24-amino acids at the cytoplasmic tail. Upon further passaging, the mutation at the S1/S2 site, as well as the stop mutation resulting in Δ24-amino acids, became dominant. Their occurrence and development, coinciding with a reduction in the VSV-G, a change in the appearance of CPE and increased viral titers suggest that these mutations may be crucial for the adaptation of rVSV-ΔG-spike to replicate in Vero E6 cells.

**Table 1:** Spike mutations acquired during serial passaging: A summary of the mutations in the spike that were acquired during serial passaging of rVSV-ΔG-spike, showing the synonymous mutation (upper row), the nonsynonymous mutation at the S1/S2 cleavage site and the nonsense mutation introducing a sto codon leading to a truncation of 24 amino acids at the cytoplasmic tail of the spike protein. *: stop codon, Ref codon: Reference codon, Alt codon: Altered codon.

| mutations | Ref codon | Alt codon |
| --- | --- | --- |
| **P507P** | **CCA** | **CCT** |
| **R685G** | **AGG** | **GGG** |
| **C1250\*** | **TGT** | **TGA** |

### Surface antigenic similarity between rVSV-ΔG-spike and SARS-CoV-2

We next evaluated the correlation between spike structures in both SARS-CoV-2 and rVSV-ΔG-spike infected cells. Such correlation may indicate an efficient and relevant immune response elicited by rVSV-ΔG-spike. Indeed, spike protein was detected by COVID-19 human convalescent sera in both rVSV-ΔG-spike and SARS-CoV-2 infected Vero-E6 cells at 24 hours post-infection (Fig. 3A). Next, we determined the ability of several COVID-19 convalescent human sera to neutralize either rVSV-ΔG-spike, or native SARS-CoV-2, in a plaque reduction neutralization test (PRNT). The NT_50_ values, the dilution at which 50% neutralization was observed, were determined for each serum sample, for neutralization of either rVSV-ΔG-spike or SARS-CoV-2. Notably, all tested human sera showed efficient neutralization of both rVSV-ΔG-spike and SARS-CoV-2. We observed a strong correlation (R^2^=0.911, p<0.0001) between the potency of human sera to neutralize rVSV-ΔG-spike and SARS-CoV-2 (Fig. 3B). Consequently, rVSV-ΔG-spike was demonstrated as an authentic surrogate for SARS-CoV-2, and thus suitable for eliciting the desired immune response.

**Figure 3:**
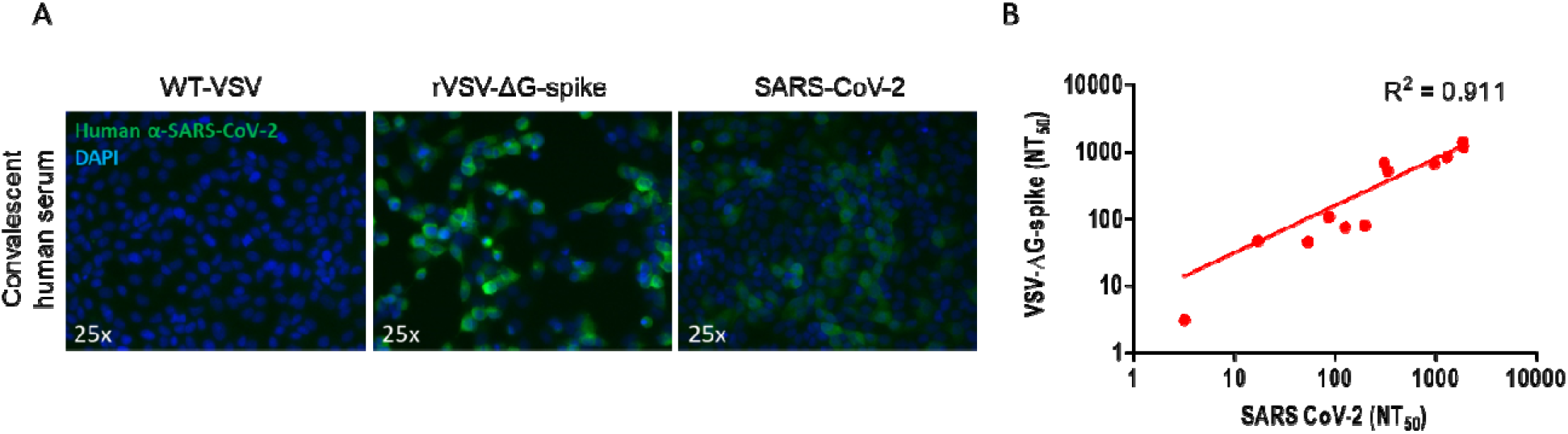
Surface antigenic similarity of rVSV-ΔG-spike and SARS-CoV-2: (A) Immunofluorescent images of Vero E6 cells infected with either WT-VSV (left panel), rVSV-ΔG-spike (middle panel), or SARS-CoV-2, stained with serum from COVID-19 human convalescent serum (right panel). (B) Correlation analysis of neutralization of rVSV-ΔG-spike and SARS-CoV-2 by a panel of sera from COVID-19 convalescent patients. For each sera (n=12), NT_50_ values were determined for neutralization of rVSV-ΔG-spike, or SARS-CoV-2. The NT_50_ values were plotted to determine the correlation between th neutralization assays. R^2^=0.911.

### rVSV-ΔG-spike vaccine efficacy in a COVID-19 hamster model

To evaluate the efficacy of rVSV-ΔG-spike as a candidate vaccine against SARS-CoV-2, we established a small animal model for COVID-19 using golden Syrian hamsters (*Mesocricetus auratus*).

In order to establish the hamster model, animals were infected with SARS-CoV-2 intranasally (i.n.) with dose of 5×10^4^, 5×10^5^ or 5×10^6^ pfu and monitored for body weight changes. As shown in Fig. 4A, animals displayed weight loss of up to 3%, 5%, and 16%, respectively, in a dose dependent manner. Therefore, a dose of 5×10^6^ pfu was determined as the inoculation dose for further experiments. Also, histological sections of lungs seven days post-infection were performed (Fig. 4B-G). Lungs of hamsters infected with 5×10^6^ pfu/animal (Fig. 4C, E, G) show focal patches of inflammation, pleural invagination and alveolar collapse, large amounts of inflammatory cells infiltration, as well as hemorrhagic areas. Edema was also observed, accompanied by protein-rich exudates. Moreover, immunostaining of the infected lung with an RBD-specific antibody showed presence of SARS-CoV-2 (Fig. 4G). Conversely, lungs extracted from naive hamsters did not show any evidence of tissue damage (Fig. 4B, D) nor stained positive for viral presence (Fig. 4F).

**Figure 4.**
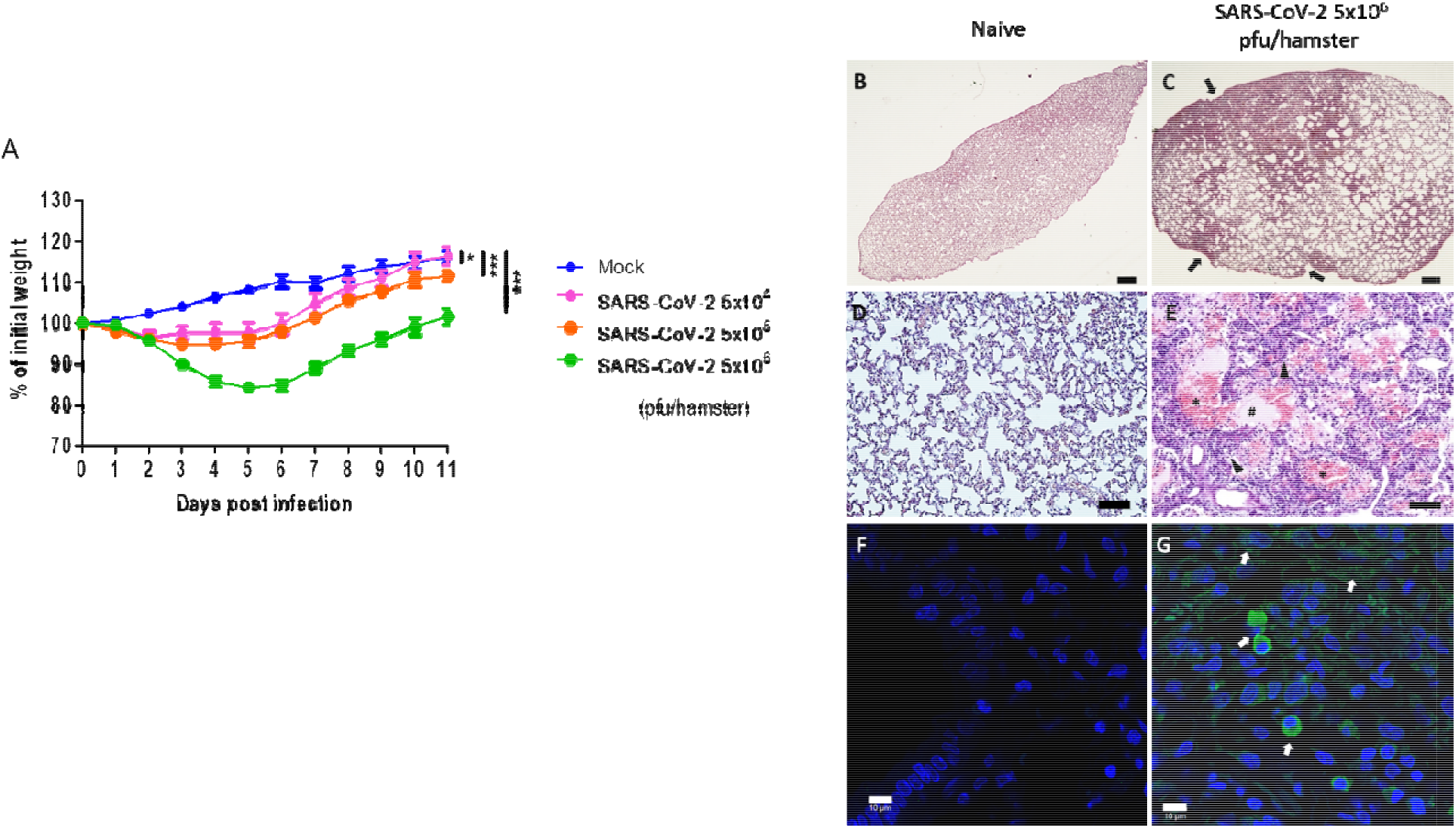
Establishment of golden Syrian hamster SARS-CoV-2 model: (A) Body weight changes of hamsters infected with SARS-CoV-2 at either 5×10^4^ (n=7), 5×10^5^ (n=7), or 5×10^6^ (n=10) pfu/hamster, compared to mock-infected hamsters (n=6). p<0.05 = *, p<0.001 = *** (B-G) General histology (H&E) an SARS-CoV-2 Immunolabeling of hamster lungs. Lungs were isolated and processed for paraffi embedding from naive (B, D, F), and SARS-CoV-2 5×10^6^ 7dpi (C, E, G) hamster groups. Sections (5μm) were taken for H&E staining (B-E) and SARS-CoV-2 immunolabeling (F-G, DAPI-Blue, SARS-CoV-2-Green). Images B-C: magnification-x2, bar= 100μm; Pictures D-E: magnification-x10, bar= 100μm; Pictures F-G: magnification-x60, bar= 10μm. Black arrows indicate patches of focal inflammation, pleural invagination and alveolar collapse. “*”-indicates hemorrhagic areas. “#”-indicates edema and protein ric exudates. Black arrow heads indicate pulmonary mononuclear cells. White arrows indicate SARS-CoV-positive immunolabeling. Naïve group: n=4, SARS-CoV-2 5×10^6^ 7dpi group: n=1.

Next, we examined the safety and efficacy of the rVSV-ΔG-spike vaccine in the hamster model. To that end, hamsters were vaccinated by intramuscular (i.m.) injection at increasing doses of rVSV-ΔG-spike: 10^4^, 10^5^, 10^6^, 10^7^ or 10^8^ pfu, and compared to mock-vaccinated hamsters. Following vaccination, animals were monitored daily for body weight changes and adverse reactions. No signs of lesion were observed at the site of injection (not shown). As seen in Fig. 5A, animals in all groups gained weight and did not show any signs of morbidity, suggesting that at the tested conditions and doses, rVSV-ΔG-spike is safe. The efficacy of the vaccine was evaluated for the ability of a single administration of each dose to elicit neutralizing antibodies against SARS-CoV-2 by PRNT. All tested vaccine doses induced a neutralization response in a dose-dependent manner (Fig. 5B). A dose of 10^6^ pfu was chosen for further evaluation. To that end, hamsters were vaccinated subcutaneously (s.c.) with 10^6^ pfu/hamster. Similarly to i.m. vaccination, the s.c. injection was found to be efficacious, as evident by induction of binding and neutralizing antibodies against SARS-CoV-2. First, hamsters’ sera were analyzed by immunofluorescent staining of Vero E6 cells infected with SARS-CoV-2. A strong signal was observed with the vaccinated hamsters’ sera, but not with sera from mock unvaccinated hamsters (Fig. 6A). Second, we evaluated the presence of neutralizing antibodies in the vaccinated hamsters’ sera. Vaccination of hamsters with rVSV-ΔG-spike generated a neutralizing antibody response, with NT_50_ value of 1:173 to SARS-CoV-2, whereas mock-vaccinated hamsters did not show any neutralizing response (Fig. 6B). Moreover, following vaccination, animals were monitored daily for body weight change and did not show any signs of morbidity (Fig. 6C), further supporting the safety of the rVSV-ΔG-spike as a potential vaccine.

To evaluate the efficacy of the rVSV-ΔG-spike vaccine to protect against SARS-CoV-2, hamsters were first vaccinated with 1×10^6^ pfu/hamster, and then challenged by intranasal (i.n.) instillation with 5×10^6^ pfu of SARS-CoV-2 per animal (four weeks post vaccination). Unvaccinated control animals were morbid, exhibiting up to 16% weight loss. In contrast, rVSV-ΔG-spike vaccinated hamsters did not show any significant signs of morbidity (Fig. 7A).

**Figure 5.**
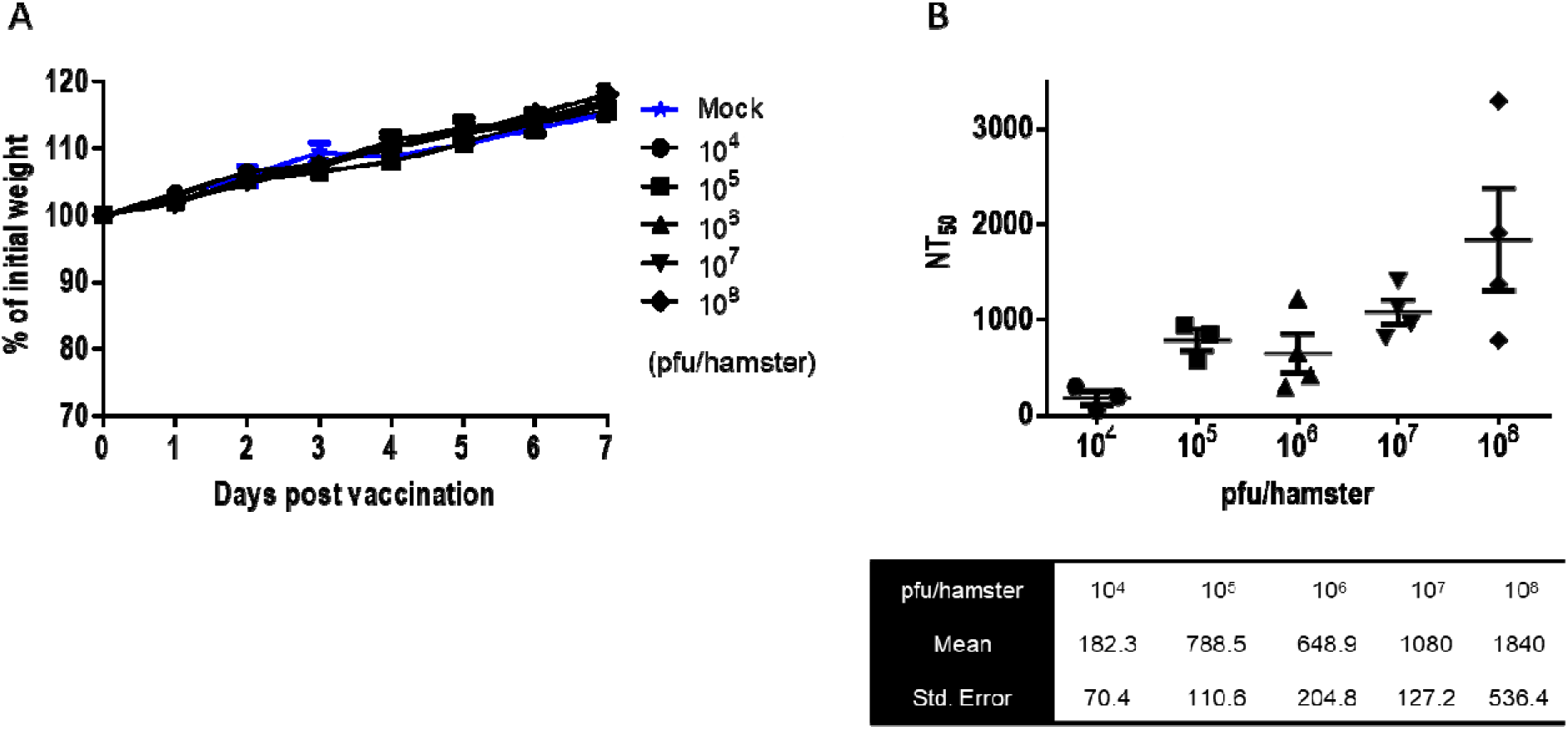
Dose-dependent vaccination of hamsters with rVSV-ΔG-spike. (A) Body weight changes of mock-vaccinated hamsters (n=4), and hamsters vaccinated with rVSV-ΔG-spike ranging from 10^4^ to 10^8^ pfu/hamster (n=8, n=10, n=10, n=10, n=8, for each vaccinated group, respectively). (B) NT_50_ values of neutralization of SARS-CoV-2 by sera from hamsters following i.m. vaccination with rVSV-ΔG-spik ranging from 10^4^ to 10^8^ pfu/hamster. n=4 for each group. Means and SEM are indicated below the graph.

**Figure 6.**
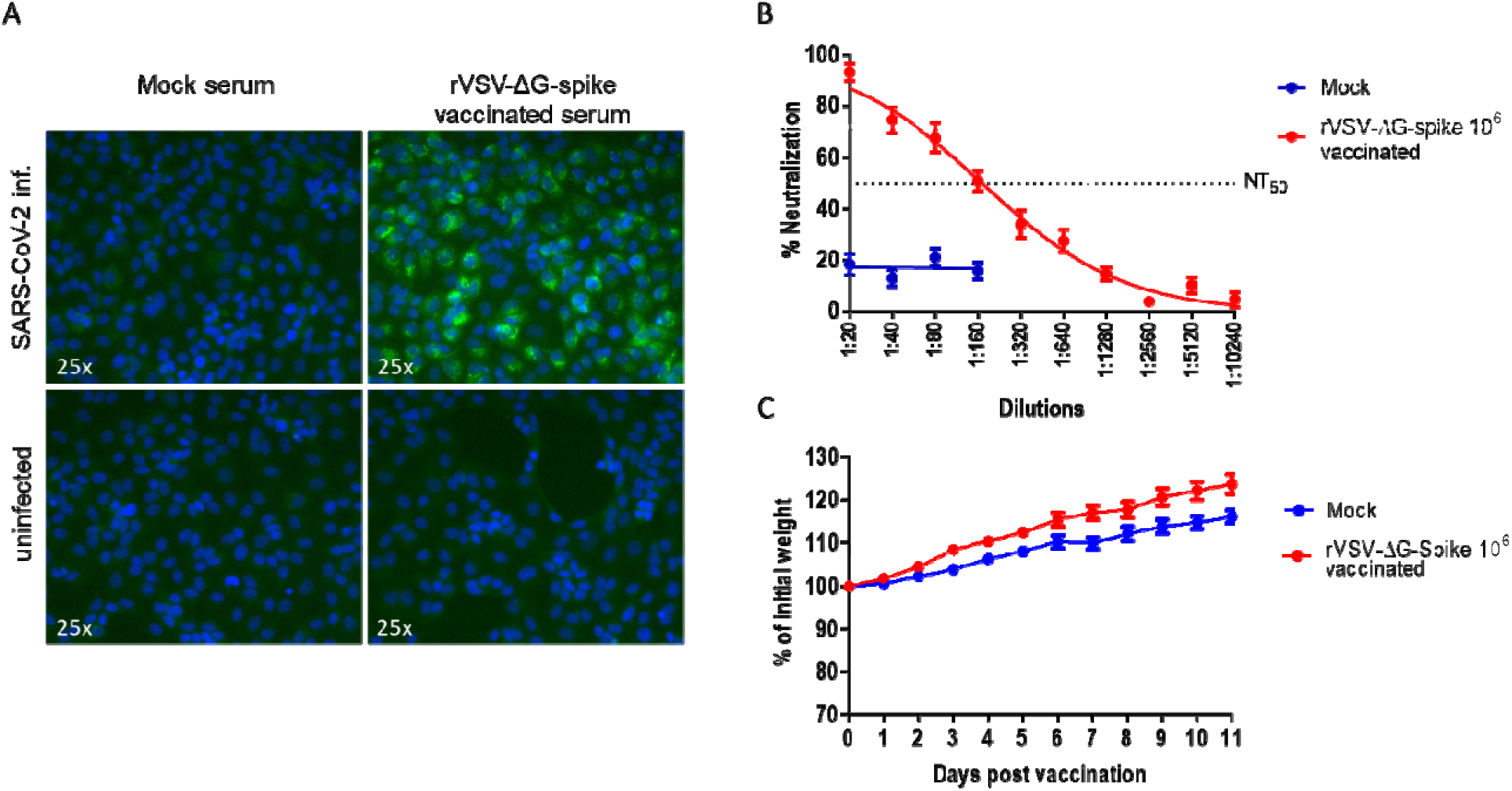
Detection and neutralization of SARS-CoV-2 by sera from hamsters following s.c. vaccinatio with rVSV-ΔG-spike: (A) Immunofluorescence images of Vero E6 cells infected with SARS-CoV-2, stai with sera from either mock-vaccinated hamsters (left panel) or rVSV-ΔG-spike vaccinated-hamsters (right panel). (B) Plaque reduction neutralization test (PRNT) of hamster sera collected from naïve hamsters (PBF, n=5) or hamsters vaccinated with rVSV-ΔG-spike 25 days following vaccination (n=5). (C) Bod weight changes of mock-vaccinated hamsters (Mock, n=16), and hamsters vaccinated with rVSV-ΔG-spike (n=8).

**Figure 7.**
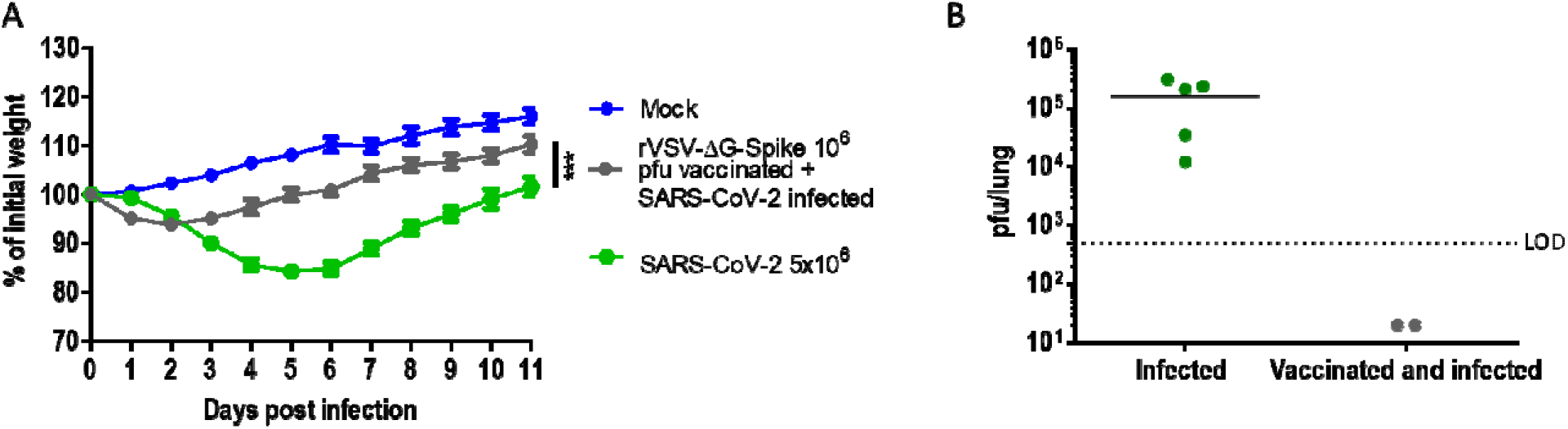
Single-dose rVSV-ΔG-spike vaccine efficacy in hamsters following SARS-CoV-2 challenge. (A) Body weight changes of hamsters infected with SARS-CoV-2 (n=12), and hamsters vaccinated wit rVSV-ΔG-spike and infected with 5×10^6^ pfu/hamster (n=10) 25 days post-vaccination, compared to mock hamsters (n=8). p<0.001 = ***. (B) Viral loads in lungs (5dpi) of hamsters infected with 5×10^6^ pfu/hamster of SARS-CoV-2 (n=5), and hamsters vaccinated with 1×10^6^ pfu/hamster of rVSV-ΔG-spike, and then infected with 5×10^6^ pfu/hamster of SARS-CoV-2 (n=2). Limit of detection (LOD, 500 pfu/lung).

Five days following challenge with SARS-CoV-2, lungs were removed and analyzed for viral load and pathological changes. Lungs extracted from infected hamsters showed average viral titers of 1.6×10^5^ pfu/lung (n=5), whereas viral titers in lungs of vaccinated and infected animals were below the limit of detection (LOD, 500 pfu/lung) (n=2) (Fig. 7B). Infected hamsters revealed extensive lung damage including focal inflammation patches, pleural invagination and massive alveolar collapse, and edema, whereas rVSV-ΔG-spike vaccinated hamsters demonstrated markedly ameliorated tissue damage (Fig. 8A-L). These results were further supported by tissue/air space analysis demonstrating significantly lower tissue/air space ratio in immunized hamsters compared to lungs of infected hamsters, but similar to that of naïve samples (Fig. 8D).

**Figure 8:**
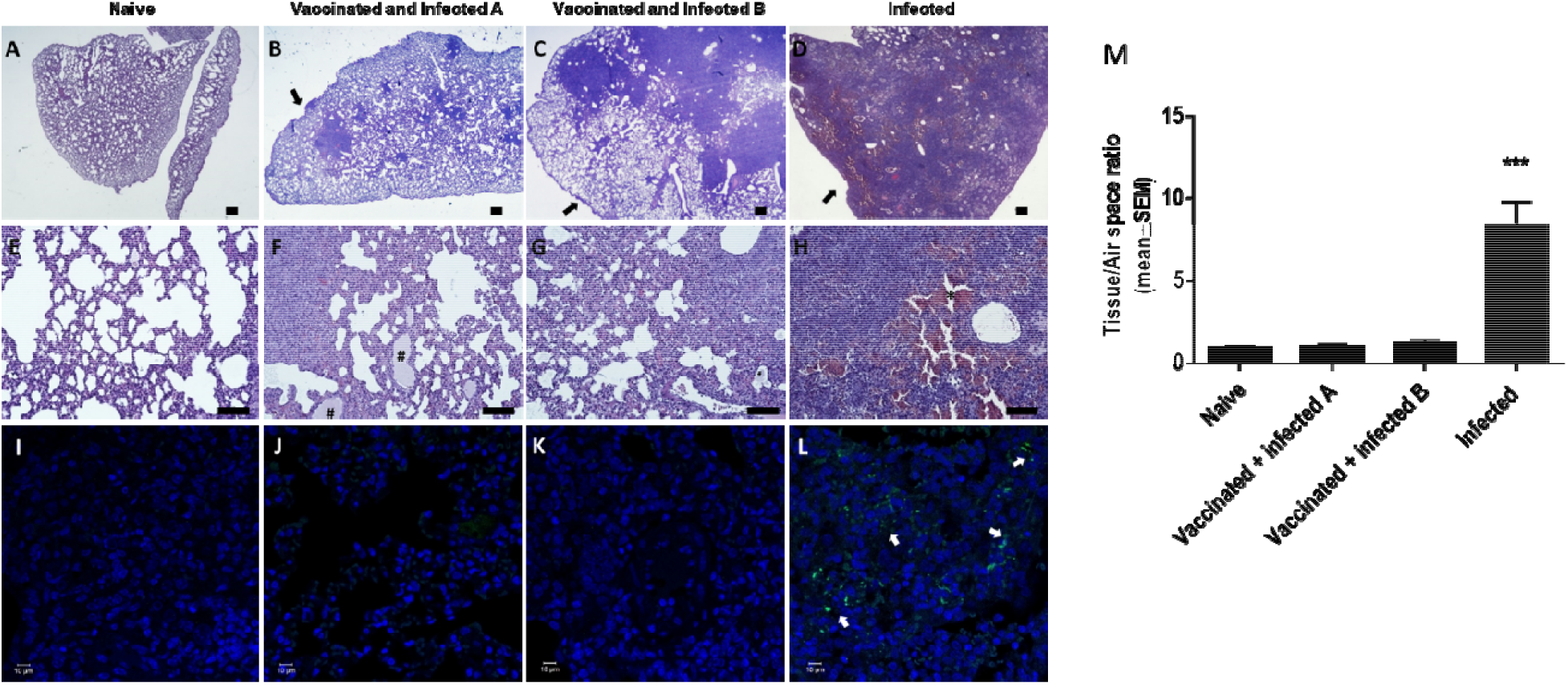
Histopathology and viral load of rVSV-ΔG-spike vaccinated and infected lungs: General histology (H&E) and SARS-CoV-2 Immunolabeling of hamster lungs with and without pre vaccination. Lungs were isolated and processed for paraffin embedding from Naïve (A, E, I), vaccinated + infected 10^6^ 5dpi (B, F, J; C, G, K), and SARS-CoV-2 5×10^6^ 5dpi (D, H, L) and groups. Sections (5μm) were taken for H&E staining (A-H) and SARS-CoV-2 immunolabeling (I-L, DAPI-Blue, SARS-CoV-2-Green). Pictures A-D: magnification-x1, bar= 100μm; Pictures E-H: magnification-x10, bar= 100μm; Pictures I-L: magnification-x60, bar= 10μm. Black arrows indicate patches of focal inflammation, pleural invaginatio and alveolar collapse. “*”-indicates hemorrhagic areas. “#”-indicates edema and protein rich exudates. Black arrow heads indicate pulmonary mononuclear cells. White arrows indicate CoV-2 positive immunolabeling. Naïve group: n=4, SARS-CoV-2 5×10^6^ pfu/animal 5dpi group: n=1, vaccinated+infecte 10^6^ pfu/animal 5dpi group: n=2. (M) Tissue/Air space ratio.

## Discussion

The urgent need for an efficacious SARS-CoV-2 vaccine led to the development of rVSV-ΔG-spike vaccine. Considering the need for induction of rapid and effective immune response, efficient production, and a strong safety profile, we chose the recombinant VSV platform. The resulting rVSV-ΔG-spike vaccine has the following attributes: 1. It efficiently replicates in cultured cells, 2. The spike is robustly expressed both on the viral surface and in infected cells, 3. It induces antibodies that effectively bind and neutralize the infectious SARS-CoV-2 virus – strongly supporting the conservation of the immunogenic spike structure. The vaccine proved effective in the golden Syrian hamster model. We show that the rVSV-ΔG-spike vaccine is safe, well tolerated, elicits antibodies, is able to bind and neutralize SARS-CoV-2 efficiently, and offers protection from high-dose SARS-CoV-2 challenge in concordance with viral clearance. Conversely, unimmunized hamsters display rapid deterioration, significant weight loss and lung damage.

The rVSV-ΔG-spike possess several features that contribute to its safety potential as a vaccine candidate. The VSV-G is considered the major virulent factor of the VSV, and its elimination is known to serve as an attenuating factor [13]. Moreover, removal of the G gene from the VSV genome, together with the expression and presentation of the spike protein, restricts the viral entry only to cells expressing the human ACE2 receptor, further significantly contributing to its safety profile [6].

Immunofluorescence and electron microscopy analysis demonstrated spike protein expression, its incorporation and proper structural organization on viral particles. Preservation of the antigenicity of the spike protein was further demonstrated by the high correlation of the neutralization capacity of a variety of human convalescent sera to neutralize both the rVSV-ΔG-spike as well as SARS-CoV-2.

The emerging pandemic calls for a rapid development of a vaccine suitable for mass production. Naturally, a platform able to achieve high titers in cultured cells is preferable. During establishment of the rVSV-ΔG-spike vaccine, the observed phenotypic changes, namely adequate spike expression and organization, reduced syncytia and increased CPE, occurred concomitantly with a gradual increase in the rVSV-ΔG-spike viral titer, reaching titers potentially suitable for human vaccination, as previously shown for rVSV-ZEBOV [13]. rVSV-ZEBOV vaccine, designed against the Ebola virus and based on the replacement of VSV-G with the Zaire ebolavirus transmembrane GP, was shown to be efficacious in clinical trials. It was approved by the FDA in December 2019 (www.fda.gov) at a dose of 2×10^7^ pfu.

During the development of the rVSV-ΔG-spike vaccine, three unique mutations emerged, including the C1250* mutation leading to a truncation of 24 amino acids at the cytoplasmic tail of the spike protein. The cytoplasmic tail of both SARS-CoV-1 and SARS-CoV-2 have been associated with the assembly of the virus and with its infectivity. This cytoplasmic tail possesses an ER retention signal [20]. Data from several studies on both SARS-CoV-1 that emerged in 2003 and the current SARS-CoV-2 reported cytoplasmic tail truncations, either deliberate or naturally occurring. These studies described the advantage of such truncations in the virus assembly, organization, and infectivity. As such, a cytoplasmic tail deletion of 19 amino-acids in spike of SARS-CoV-1 allowed efficient assembly and incorporation of the spike into VSV particles, resulting in high titers of VSV-SARS-St19 [21, 22]. A similar deletion of 19-amino acids in SARS-CoV-2 spike was also shown to enhance expression of SARS-CoV-2 in mammalian cells [23]. Moreover, several recent studies establishing SARS-CoV-2 BSL-2 platforms reported the occurrence of one or two independent stop mutations, leading to 21- or 24- amino acids truncations in the cytoplasmic tail of SARS-CoV-2 [24, 25].

Further dissection of the characteristics of the spike’s cytoplasmic tail and the effects on the SARS-CoV-1 by performing single mutations, as well as truncation mutations, in the spike [26], suggested an advantage for such mutations to cell surface expression, as well as an effect on the fusion capacity of the virus in some of the truncated forms; a 26-amino acid truncation reduced syncytia size by average of 22%, whereas a 17-amino acid truncation increased syncytia size by average of 43%, thus increasing fusion capacity of the truncated virus. The authors suggested that the 17 amino acid residues at the cytoplasmic tail exert a negative effect on spike-mediated cell fusion, and the 26-amino acids of the cytoplasmic tail domain, comprising a cysteine-rich motif, have a secondary effect on cell fusion. These cytoplasmic tail alterations creating truncations at 17- and 26-amino acids, span the 24-amino acids truncated area reported herein. Accordingly, the rVSV-ΔG-spike lacking 24-amino acids described here indeed reduced syncytia formation while maintaining high expression and presentation of the spike protein.

The nonsynonymous mutation R685G generated during passaging of the rVSV-ΔG-spike, led to a change in the S1/S2 motif RRAR, creating a RRAG site. It was shown that the S1/S2 site of the spike protein contains a multibasic motif which is processed by the host cell furin protease. This site is essential for cell-cell fusion, and was shown to mediate infection of human lung cells [7]. Additional study [27] demonstrated that a variant of SARS-CoV-2 with a deletion in the furin site has reduced virulence to hamsters, further substantiating the important role of this RRAR motif in virus infection and pathogenesis. This suggests that the R685G mutation in the rVSV-ΔG-spike may contribute to its attenuation.

Taken together, our data suggests that both the stop mutation leading to Δ24, and the S1/S2 cleavage site mutation of the rVSV-ΔG-spike vaccine candidate contribute to spike protein expression, efficient assembly of the viral particle and its ability to replicate and propagate in Vero E6 cells, thus improving its stability, attenuating its virulence and contributing to the overall safety of this candidate vaccine.

Previous works presented the golden Syrian hamster as a model for SARS-CoV-2 pathogenesis and transmission [28, 29]. In this work, we implemented the golden Syrian hamster model to evaluate the rVSV-ΔG-spike vaccine safety and efficacy. The hamster model is a robust and reproducible model as evident by the dose-dependent response in body weight to different SARS-CoV-2 infection doses. Also, vaccination of the hamsters 25 days pre-exposure to SARS-CoV-2 resulted in a dose-dependent induction of neutralizing antibodies against SARS-CoV-2, thus providing a reliable tool for estimating vaccination efficacy. Vaccination of hamsters with rVSV-ΔG-spike prior to inoculation with SARS-CoV-2 provided protection, as demonstrated by minimal weight loss, followed by a remarkable recovery within a few days and a complete return to baseline by day 5. This is contrasted with the unvaccinated hamsters, displaying a prolonged disease, peaking at day 5, followed by a slow and gradual recovery. Importantly, analysis of lungs extracted from vaccinated animals showed minimal pathology and no detectable virus opposed to SARS-CoV-2 infected hamsters showing extensive damage and high viral loads.

In conclusion, we generated rVSV-ΔG-spike, a recombinant replication-competent VSV-based vaccine candidate expressing the SARS-CoV-2 spike protein. The rVSV-ΔG-spike resembles the SARS-CoV-2 in spike expression properties, antigenicity, and ability to induce neutralizing antibody production. Moreover, single-dose vaccination of hamsters with rVSV-ΔG-spike elicits a safe, effective, and sufficient neutralizing antibody response against SARS-CoV-2 challenge. The vaccination provided protection against SARS-CoV-2 inoculation, as manifested in the rapid return to normal physiological parameters lung protection and rapid viral clearance. These results pave the way for further examination of rVSV-ΔG-spike in clinical trials as a vaccine against SARS-CoV-2.

## Author Contributions

Y.Y.R., H.T., S.M., B.P., H.A., E.B.V., L.C., Y.V., A.B.S., E.B.D., A.S., H.L., N.P. and T.I. conceived and designed the experiments. Y.Y.R, H.T., S.M., B.P., H.A., E.B.V., A.S., A.B.D., N.P. and T.I. analyzed the data. Y.Y.R., H.T., S.M., O.S., I.G., N.P. and T.I. wrote the manuscript. N.E., R.Z., E.M., O.M., A.B.D., N.P. and T.I. contributed in conceptualization. O.S., O.I., D.S., I.C.G., A.Z. and A.B.D. contributed in bioinformatics. O.S., L.C., S.L., S.W., R.A., R.P., L.C., and E.L. contributed in technical support. E.M., H.G., S.L. and O.L. contributed in histology and electron microscopy. S.Y. and S.C.S. added fruitful discussions. N.P. and T.I. supervised the project. All authors commented on and approved the manuscript.

## Acknowledgments

We acknowledge the IIBR administrative personnel for their commitment to the project. We thank Amir Rosner, Beni Shareabi and Yossi Schlomovitch for animal husbandry. We thank Eilat Shinar for serum samples of COVID-19 convalescent patients. We thank Eran Bacharach for protocols and reagents.

